# Immune reactivity to neurofilaments and dipeptide repeats in ALS progression

**DOI:** 10.1101/2020.02.25.965236

**Authors:** Fabiola Puentes, Vittoria Lombardi, Ching-Hua Lu, Ozlem Yildiz, Angray Kang, Ahuva Nissim, Pietro Fratta, Adrian Isaacs, Yoana Bobeva, Andrea Malaspina

## Abstract

**Objective:** To test antibody response and formation of immune-complexes to neurofilaments and dipeptide-repeats, the translational products of the mutated C9orf72 gene, as potential biomarkers for clinical stratification of amyotrophic lateral sclerosis (ALS).

**Methods:** Using neurofilament protein isoforms plasma expression as reference, antibodies and immune-complexes against neurofilament-light, medium and heavy chain and poly-(GP)-GR dipeptide-repeats were tested in blood from 105 fast and slow progressing ALS patients, 26 C9orf72 mutation carriers (C9+ve) ALS patients and 77 healthy controls (HC) using single-molecule and immune-capture assays. Longitudinal antibody/immune-complex responses were measured in serial blood samples from 37 (including 11 C9+ve) patients.

**Results:** Antibodies and immune-complex reactivity was higher in ALS patients than HC, particularly in C9+ve ALS patients, and modestly correlated with total neurofilament protein isoforms (r:0.24 *p*=0.002; r:0.18 *p*=0.02 respectively). Neurofilament-light immune-complexes and neurofilament-heavy antibodies had the best diagnostic performances distinguishing ALS subtypes from HC (AUC=0.68 *p*<0.01; AUC=0.68 *p*<0.001 respectively). Neurofilament-light immune-complexes (AUC=0.69 *p*<0.01) and poly-(GP) dipeptide-repeats antibodies (AUC=0.71 *p*<0.001) separated C9+ve from C9-ve patients. Multivariate mortality hazard ratio and Kaplan-Meier analyses showed low neurofilament-heavy antibody levels associated with increased survival. Longitudinal analysis identified raising levels of antibodies against neurofilaments in fast progressing ALS and of neurofilament-light immune-complexes in C9+ve patients.

**Interpretation:** C9+ve and fast progressing ALS patients have a distinct neurofilament and dipeptide-repeat immuno-phenotype, with increasing levels of blood neurofilament-light immune-complexes and neurofilament antibodies with disease progression. The study of the expression of these biomarkers in the natural history of ALS may shed light on disease initiation and progression and provide novel pharmacodynamic biomarkers in emerging C9orf72 gene silencing therapies.

## Introduction

The lack of biomarkers supporting diagnosis, prognosis and an accurate definition of treatment response impact on the development of a comprehensive therapeutic strategy for amyotrophic lateral sclerosis (ALS), a rapidly progressing and fatal neurodegenerative disorder^1^. The genetic mutations causing ALS may partially explain this conditions’ clinical heterogeneity, where survival from disease onset varies from less than a year to more than a decade^1-3^. The pathology underpinning the clinical heterogeneity of ALS is centred around the role of disordered proteins and on the modifying effect on progressive neuronal loss of disease-specific immune responses, such as the reduction of T-regulatory cells predominantly in fast progressing ALS, which is likely to induce a state of diminished self-immune tolerance and the formation of antibodies (Abs) and immuno-complexes (ICs) to brain proteins^4,5^. Intronic expansions of the C9orf72 gene, the most common genetic mutation in ALS, are also thought to enhance a state of autoimmunity possibly induced by loss of function of translation products like dipeptide repeats (DPR)^6^.

The detection in cerebrospinal fluid (CSF) and blood of brain proteins like neurofilaments (Nf) is thus far the most informative neurochemical marker of ALS^7,8^. Nf homeostasis in biofluids and the stoichiometry of Nf isoforms light (Nf-L), medium (Nf-M) and heavy (Nf-H) depend on rate of axonal loss and on the antigen-clearing effect of the immune system, represented by the formation of antibodies/immune-compIexes against Nf (Nf Abs; Nf ICs) and on Nf compartmentalization into circulating heterogeneous macromolecular aggregates^9^. Nf-L Abs levels in blood and Abs against the oxidized form of the translation products of the gene superoxide dismutase 1 (oxSOD1 Abs) have already been investigated as biomarkers of ALS. Nf-L Abs reach a peak in blood at a late stage of ALS and correlate with functional impairment and survival, while oxSOD1 Abs are over-expressed in blood from slow progressing ALS^10,11^. These Abs may potentially exert a modifying effect on disease progression.

Moreover, the formation of Nf ICs may skew Nf detection, as endogenous Abs would compete with immunoassays Abs for epitope recognition. This may explain why a sensitive assay for poly (GP) DPR detection is only available in CSF, where antibody levels concentration is much lower compared to blood^12^. Whilst poly (GP) DPR detection in CSF remains the most obvious pharmacodynamic biomarker for ongoing antisense oligonucleotide (ASO) trials in C9+ve ALS, serial lumbar punctures to monitor treatment response in advanced ALS patients may be impractical. In contrast, detection of stable molecules like Abs/ICs in blood would be less invasive, less affected by sample quality and relying on relatively inexpensive Abs/ICs capture methods in a small sample size (1-2 µl).

Using Nf antigen levels as reference, here we investigate the humoral response against Nf and DPR as biomarker of disease progression in ALS. We show that Abs/ICs against specific Nf and DPR isoforms provide novel molecular classifiers for the clinical stratification of ALS. The panel of immunological markers proposed here may complement Nf and poly (GP) analysis in the phenotypic and genotypic characterization of ALS, in the prediction of survival and prognosis.

## Patients and Methods

### Subjects

Shortly after diagnosis (average 2 months (±2)) and clinical review by an ALS neurologist, patients fulfilling the El Escorial Criteria for probable, laboratory supported, probable and definite ALS^13^ were consented along with age and sex matched healthy controls. Written informed consent was obtained from all participants. Ethical approval for inclusion in the ALS biomarkers study was obtained from the East London Research Ethics Committee (REC reference 09/H0703/27). Exclusion criteria for both ALS and HC included comorbidities likely to change Nf expression and immune response, including history of brain or spinal cord inflammatory, neurodegenerative disorders and injury, autoimmune disorders, recent treatment with steroids, immunosuppressants and immunoglobulins.

All participants were sampled at baseline after diagnosis for cross-sectional analyses, while a subset of individuals was sampled serially to allow for longitudinal biomarkers studies.

The ALS functional rating scale revised (ALSFRS-R) score was recorded at each visit (48 is equivalent to healthy state, the lower the score the higher the neurological disability), while rate of disease progression was defined as progression rate to last visit (PRL; ALSFRS-R at onset assumed at 48, minus ALSFRS-R at the time of sampling, divided by the time interval between onset and sampling in months)^8^. Genetic analysis for the C9orf72 mutation and ATX2 variant was undertaken for all ALS cases.

### Plasma extraction

Blood samples were drawn by venepuncture and collected in anticoagulant (EDTA) coated tubes (BD vacutainer, UK), centrifuged at 3500 rpm for 10 minutes at room temperature. Plasma was aliquoted in polypropylene tubes (Nunc, UK) and stored at −80°C until use.

### Proteins and Peptides

For Abs detection, the following capture proteins were selected and coated onto ELISA plates: 1) human-recombinant Nf-L (MyBiosource, USA) and Bovine Nf-L, Nf-M, Nf-H (Progen, Germany) for Nf and 2) 20 amino acid-long biotinylated peptides (10 × GA, GP and GR) for DPR ^14,15^. The Poly(GA)10, poly(GP)10 and poly(GR)10 peptide repeats were synthesized at the National Physical Laboratory UK and biotinylated at the amino terminal region with purity >90%.

### Quantitative Antibody determination

Plasma samples were tested for antibody detection by enzyme-linked immunosorbent assay (ELISA) as previously reported^16^. Nunc-Immuno microtiter 96-well solid plates (Thermo Fisher Scientific, UK) were coated with 2 μg/ml recombinant human Nf-L (MyBiosource, USA) or 3 μg/ml Bovine Nf-L, Nf-M or Nf-H (Progen, Germany) in carbonate buffer pH 9.6, overnight at 4°C. Plates were blocked with 2% bovine serum albumin PBS/BSA and plasma samples were assessed at 1:100 dilution in triplicates. After one-hour incubation, antibody binding was detected with horseradish-peroxidase (HRP)-conjugated goat anti-human IgG (Sigma, UK), used at 1:5000 dilution. The reaction was developed with TMB (3,3′,5,5′-tetramethyl-benzidine substrate, Thermo Fisher Scientific, UK) and stopped by the addition of 2 M hydrochloric acid (Fluka, UK).

NeutrAvidin plates (Thermo Scientific, UK) were incubated with 10 μg/ml biotinylated poly (GP), poly (GR) and poly (GA) DPR peptides in Tris-buffered saline (25mM Tris, 150mM NaCl pH 7.2, 0.1% BSA, 0.05% Tween-20) followed by incubation with plasma and washing with TBS. Bound IgG were detected by addition of enzyme-labelled secondary antibodies as described above. The reaction was measured in a Synergy HT microplate reader (Bio-Tek instruments, VT).

### Immune complexes determination in plasma

Quantification of Nf and DPR immune-complexes (ICs) was carried out by ELISA as previously reported^17^. Microtitre 96-well plates (Thermo Fisher Scientific, UK) were coated overnight at 4°C with 0.1 mL of 0.5 µg/ml rabbit polyclonal anti-human Nf-L (Abbexa, UK), anti-Nf-M (Abbexa, UK), anti-Nf-H (Novus Biological, UK), or polyclonal rabbit anti-(GP) and anti-(GR) antibodies (kindly provided by Professor Adrian Isaac, UCL, UK). All antibodies were diluted in carbonate buffer pH 9.6. After blocking with PBS/2% BSA, plates were incubated with plasma samples at 1:100 dilution for 1 h at room temperature. After washing with PBS 0.1% Tween-20, (HRP)-conjugated goat anti-human IgG (Sigma, UK) was added and incubated for 1 h at room temperature. The reaction was developed with TMB and measured at 450 nm.

### Measurement of plasma Nf-L and Nf-H levels

Analysis of Nf-L and Nf-H protein expression in human plasma was undertaken by single molecule array (Simoa) technology using a digital immunoassay HD-1 Analyzer (Quanterix, Lexington, MA). Standards, primary and secondary antibodies, detection range including lower and upper limits of detection were specified in the manufacturer’s conditions of the commercial assays. Analysis of Nf-M protein isoform was not attempted as currently available immunoassays have not been extensively tested in ALS ^18^.

Genotyping of C9orf and ATAX-2 genes in the ALS population in this study was performed as previously reported^19^.

### Statistical Analysis

Continuous variables were expressed as median (interquartile range; IQR); non-parametric group analysis was performed using Kruskal–Wallis one-way test of variance on ranks and after log-transformation of raw values. Spearman’s correlation was used to evaluate the relation between Abs, ICs expression and ALSFRS-R and PRL. To investigate changes in longitudinal plasma analytes, Nf Abs and ICs plasma levels slopes were calculated between the late time point and baseline measurements and expressed as ratio of time intervals between the same time points. These ratios were further analysed in relation to changes in ALSFRS-R with the disease progression. In addition, Nf Abs and ICs levels were interpolated with the z-score, the average change of the analytes in all time points considered, in both ALS-F and ALS-S. Correlation analysis (Spearman’s) was also used to test associations between analytes expression and functional scores and between the Abs/ICs slopes and ALSFRS-R slopes to evaluate patters of disease progression.

Receiving operator curve (ROC) non-parametric analyses and their corresponding 95% confidence intervals (CI) were calculated to evaluate sensitivity and specificity for the discrimination of patient subgroups and separation from healthy controls, based on Abs and ICs levels against Nf and poly-DPR isoforms.

Kaplan-Meier curves and log rank test were performed to define survival across groups, along with Cox proportional hazards models with multivariate analysis, adjusted for sex, age, onset-diagnosis time interval, ALSFRS-R scores, site of onset (bulbar, upper limbs region, lower limbs region and respiratory/thoracic region). A *p* value of less than 0.05 was considered statistically significant. Data were analysed using Prism (Version 8.0, GraphPad Software, San Diego, CA, USA).

## Results

### Study population

Demographic and clinical characteristics for the cross-sectional and longitudinal cohorts are summarized in Tables 1 and 2 respectively. These included 105 ALS patients comprising 42 with fast disease progression (ALS-F; PRL > 1.0), 37 slow progressing cases (ALS-S; PRL < 0.5), 26 ALS patients carrying the C9orf72 hexanucleotide repeat expansion (C9+ve), 3 C9orf72-ve (C9-ve) patients found to have the ATX-2 gene expansion and 77 healthy controls (HC). The median age at baseline was 56.7 years in HC and 65.8 in ALS patients. For the cross-sectional cohort, the median and IQR of time since disease onset was 3.14 (2.49 – 6.19 years) in ALS-S, 0.97 (0.66 – 1.60 years) in ALS-F and 1.51 (1.05 – 2.19 years) in C9+ve ALS. The female/male gender ratio was 0.94 in ALS-S, 1.4 in ALS-F, 1 in C9+ve and 1.8 in HC (Table 1).

**Table 1.** Cross-sectional study. Demographic and clinical features of the study population.

| Cross-sectional data | ALS (Slow progression) | ALS (Fast progression) | C9orf72 positive | Healthy subjects |
| --- | --- | --- | --- | --- |
| Number (n) | 37 | 42 | 26 | 77 |
| Gender Female/Male | 12/25 | 25/17 | 13/13 | 49/27 |
| Age at baseline (years)<sup>a</sup> | 66.70 ± 12.33 | 64.92 ± 11.25 | 56.77 ± 10.30 | 58.56 ± 8.55 |
| **Site of disease onset:** |  |  |  |  |
| Bulbar | 7 | 17 | 8 | NA |
| Limb | 28 | 19 | 14 |  |
| Bulbar and Limb | 1 | 3 | 2 |  |
| Other/Unknown | 1 | 3 | 2 |  |
| Years since onset of weakness (Disease duration)<sup>b</sup> | 3.14<br>(2.49 – 6.19) | 0.97<br>(0.66 – 1.60) | 1.51<br>(1.05 – 2.19) | NA |
| Years since diagnosis<sup>b</sup> |  |  |  | NA |
| Baseline ALSFRS-R<sup>a</sup> | 38.73 ± 7.59 | 32.90 ± 7.57 | 35.38 ± 8.88 | NA |
| Survival time (month)<sup>b</sup> | 42.00<br>(37.00 – 44.00) | 23.26<br>(15.93 – 29.79) | 28.78<br>(19.09 – 37.17) | NA |
| PRL<sup>a</sup> | 0.22 ± 0.13 | 1.65 ± 0.95 | 0.89 ± 0.58 | NA |
| % Caucasian / no-Caucasian /no data | 97.3/2.7/0 | 88.1/11.9/0 | 92.3/7.7/0 | 55.85 / 5.19 / 38.96 |

<sup>a</sup>Data are presented as mean ± standard deviation (±SD).
<sup>b</sup>Data are presented as median and interquartile range (IQR)
(ALSFRS-R): Amyotrophic lateral sclerosis Functional Rating Scale Revised
PRL: Progression rate at last visit
NA, not applicable

**Table 2.**
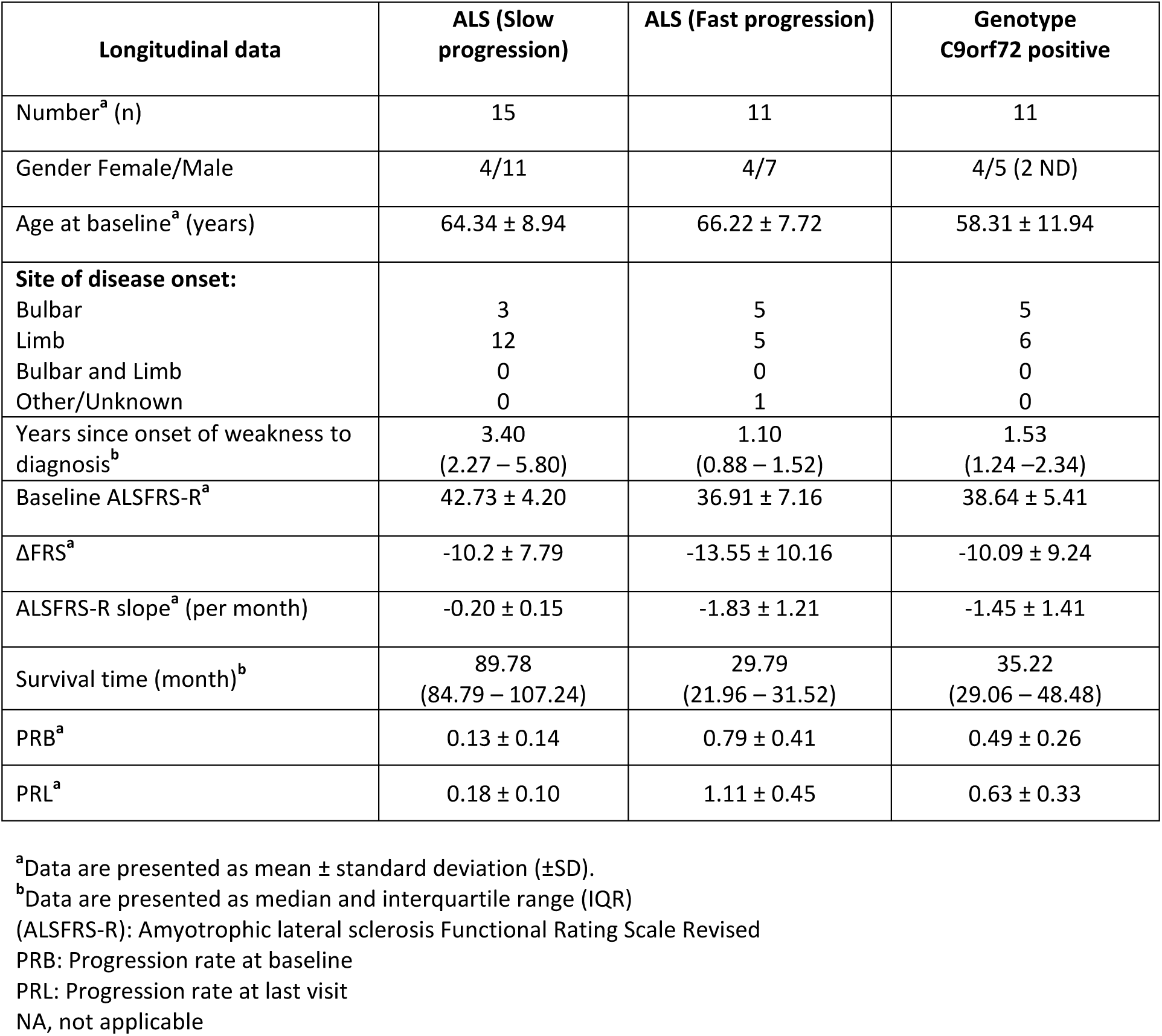
Demographic and clinical features of the Longitudinal study population.

15 ALS-S, 11 ALS-F and 11 C9+ve patients (ALS-F and ALS-S) had further analyses in blood samples drawn serially in up to 4 distinct time points. Follow-up visits occurred on average in 3 to 8 months intervals. 58.1% of patients in the cross-sectional study and 62.1% in the longitudinal study had limb onset disease; the rest had predominantly bulbar clinical features at presentation. The average baseline ALSFRS-R score was 37.4±16.8 for the cross-sectional cohort and 40.3±6.2 for the longitudinal cohort (Tables 1 and 2).

### Abs and ICs assay performance

Accuracy of immunoassays for both Abs and ICs was improved by the inclusion of standard curves. Plates were pre-coated with anti-human IgG (Fc specific) antibody (Sigma, UK) to capture purified human IgG (Sigma, UK) which was serially diluted (1:2) from 375 to 0.02 ng/ml to generate a standard calibration curve. After incubation with peroxidase-conjugated goat anti-human IgG (whole molecule), TMB substrate was added for detection. Optical densities were normalised by subtracting the absorbance derived from uncoated wells. Plasma Abs and the proportional amount of IgG and ICs (pg/ml) were quantified by interpolation of optical densities from the standard curve.

For each round of Abs and ICs ELISA, plasma samples from ALS patients and healthy controls were equally distributed on each plate and measured in duplicate. The intra-assay coefficients of variation (%CVs) were 9.3%, 9.9%, 11.2%, 6.8% and 5.0% for detection of Nf-L, Nf-M, Nf-H, GP and GR antibodies respectively. The inter-assay %CVs were 18.1%, 9.9%, 14.6%, 10.4% and 13.4% for detection of Nf-L, Nf-M, Nf-H, GP and GR antibodies respectively. For immune-complexes the intra-assay %CVs were 11.2%, 10.6%, 8.7%, 11.1% and 10.0% for Nf-L, Nf-M, Nf-H, GP and GR respectively. The inter-assay %CVs were: 16.0%, 14.3%, 12.7%, 10.3% and 10.8% for Nf-L, Nf-M, Nf-H, GP and GR immune-complexes respectively. For the Abs assays, the lower limit of detection (LLD) across experiments was 0.058 ng/ml and the upper limit of detection was 280 ng/ml, while for ICs, it was 0.03 ng/ml and 65 ng/ml respectively. Missing data (i.e. technical replicates below the lower limit LLD) for Abs was 2.3% of total measurement while for ICs was 3.2% of total measurement.

### Total neurofilaments (Nf) and dipeptide repeats (poly-DPR) immune response in ALS

We selected bovine Nf proteins (Nf-L 68 KDa, Nf-M 160 KDa and Nf-H 200 KDa; Progen), highly homologous to human Nf proteins (a single band at the expected molecular weight on electrophoresis) and with >98% purity. To account for the expected Abs cross-reactivity to different Nf chains (due to amino acid similarities across the three standard proteins), we tested both individual response and (total) reactivity to all Nf isoforms.

Total Nf (Nf-H plus Nf-L) protein isoforms expression levels in blood, total Nf (Nf-L, Nf-M and Nf-H) Abs, ICs and DPR Abs and ICs (poly (GP) and (GR)) were examined in the same plasma samples from ALS and HC (Fig.1 A1-A5). In Fig.1 B1-B5, the expression of the same markers is shown across the ALS-S, ALS-F and C9+ve variants of ALS.

**Figure 1.**
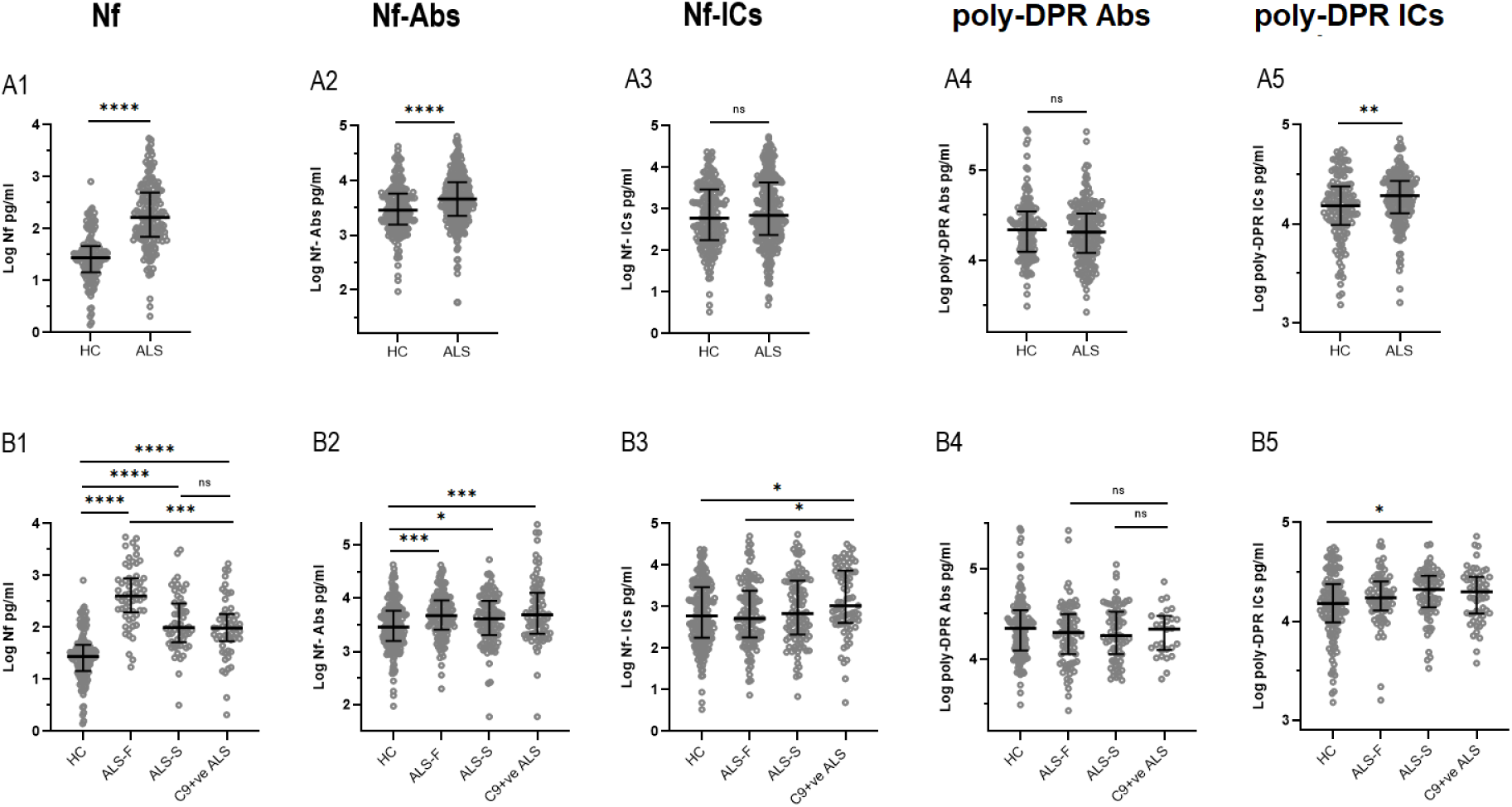
Immune response to neurofilaments (Nf) and dipeptide repeats (DPR) peptides in blood from patients with amyotrophic lateral sclerosis (ALS) and healthy controls (HC). (**A1**): the combined plasma expression of neurofilament light and heavy chain (Nf-L and Nf-H) Nf isoforms. (**A2**): Abs against Nf-L, Nf-M and Nf-H (Nf-Abs). (**A3**): Nf-L, Nf-M and Nf-H immuno-complexes (Nf-ICs). (**A4)**: Abs against poly (GP) and poly (GR) variants of dipeptide repeats (poly-DPR Abs) and (**A5**) poly (GP) and poly (GR) variants immuno-complexes (poly-DPR ICs). Levels of Nf and Nf-Abs are significantly higher in the ALS cohort compared to HC (**A1, A2**, *p*<0.0001). No significant differences in Nf-ICs and poly-DPR Abs levels are observed between ALS and HC (**A3, A4**) while plasma poly-DPR ICs are higher in ALS patients compared to HC (**A5**, *p*<0.01). Analysis of neurofilament isoforms (Nf), antibodies (Abs) and immuno-complexes (ICs) against Nf and dipeptide repeats (DPR) variants in fast progressing ALS (ALS-F), slow progressing ALS (ALS-S) and ALS individuals with a known C9orf72 (C9+ve) or ATX-2 (ATX+ve) genetic mutation and HC. (**B1):** higher expression levels for Nf in ALS-F compared to HC (*p*<0.0001) and C9+ve ALS (*p*<0.001). Nf is also significantly overexpressed in ALS-S and C9+ve ALS compared to HC (*p*<0.0001). (**B2):** Nf-AbS plasma expression in C9+ve ALS and ALS-F is higher compared to HC and ALS-S (*p*<0.001). (**B3**): higher expression of Nf-ICs in C9+ve ALS compared to HC (*p*<0.05) and ALS-F (*p*<0.05). (**B5):** poly-DPR ICs are significantly increased in ALS-S compared to HC (*p*≤0.05).

Nf and matched Abs were over-expressed in blood from ALS individuals compared to HC (A1, A2; *p*<0.0001). No difference in the total level of Nf-ICs was observed in ALS patients compared to HC (A3). ICs and not Abs against DPR were up-regulated in ALS blood compared to HC (A5, *p*=0.0029).

In phenotypic and genetic variants of ALS, total Nf appeared more expressed in blood from ALS-F compared to HC (Fig.1 B1, *p*<0.0001), similarly to matched Abs and ICs against Nf, with C9+ve ALS and ALS-F showing the highest expression levels (Fig.1 B2, *p*=0.0009 and B3, *p*=0.011). C9+ve patients showed higher levels of Nf IC than the ALS-F group (Fig.1 B3, *p*=0.022). No difference of poly-DPR Abs expression between ALS subgroups and HC were observed (total poly-DPR Abs, Fig.1), while poly-DPR ICs were more expressed in ALS-S compared to HC (Fig.1 B5, *p*= 0.02).

### Nf and poly-DPR isoform-specific immune response in ALS

Group analysis and diagnostic performance using ROC were carried out to examine cross-sectionally how Abs and ICs discriminate ALS patient phenotypic variants from HC, compared to Nf-L and Nf-H proteins measured in the same samples. We have previously reported no differential regulation of Nf-M plasma between ALS and HC, using a non-validated commercial assay^18^. Nf-L and Nf-H proteins were over-expressed in plasma samples from ALS patients compared to HC (*p*<0.0001) ^8,20^. Nf-L plasma expression was also significantly different in ALS-F compared to ALS-S (Fig.2 A1, *p*=0.0014). Blood expression of Nf-L and Nf-H protein isoforms in C9+ve patients were significantly reduced compared to the ALS-F subgroup and HC (Fig.2 A1, A5 *p*=0.002 and *p*=0.027 respectively). ROC analyses indicated that Nf-L and Nf-H blood proteins separated well HC from ALS patients (AUC=0.93 and AUC=0.87, *p*<0.0001 for Nf-L and Nf-H respectively), while only Nf-L blood expression discriminated between ALS-F and ALS-S (AUC= 0.89, *p*<0.0001; sensitivity 92.3% and specificity 92.5%, Fig.2 A2).

**Figure 2.**
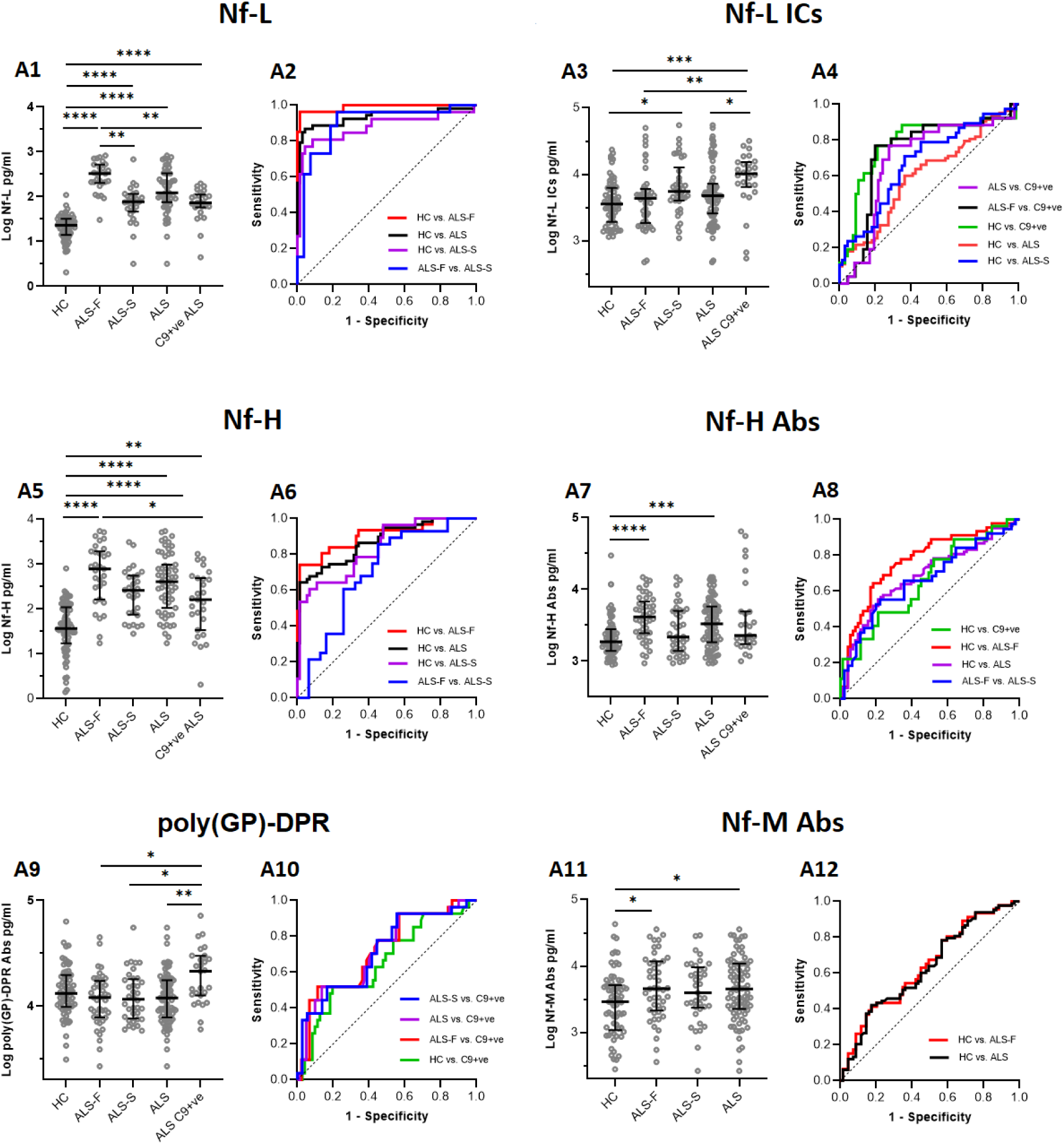
Immune response to Nf isoform proteins and to poly-Gp dipeptide repeats (DPR) in phenotypic variants of ALS and in HC. Group analysis comparing Nf-L and Nf-H protein isoforms expression, Nf-L, Nf-M and Nf-H antibodies (Abs) and immuno-complexes (ICs) expression in healthy individuals (HC), phenotypic (ALS-F, ALS-S) and genotypic (C9+ve; ATX-2+ve) variants of ALS. Receiver operating characteristic (ROC) non-parametric analysis is used to evaluate the ability of the analytes to discriminate ALS phenotypic subgroups from HC (plots indicate sensitivity against 1-specificity). Kruskal-Wallis One-way analysis was used for multiple comparisons. The scatter dot plots show the median with interquartile range. The statistical difference between groups is shown as (****p<0.0001), (***p<0.001), (**p<0.01), (p*≤0.05) or non-significant (n.s). (**A1, A2)**: Significantly higher Nf-L protein levels were observed in all ALS subgroups compared to HC (p<0.0001). ROC analysis showed that Nf-L proteins strongly discriminate ALS patients from HC and ALS-F from ALS-S (**A2**; AUC=0.93 p<0.0001 and AUC=0.89, p<0.0001 respectively). (**A5, A6**): Nf-H protein levels are increased in ALS patients and particularly in ALS-F similarly to what seen for Nf-L. (**A3, A4**): Nf-L ICs levels separate ALS-S from HC (p<0.05) and are significantly higher in C9+ve ALS patients compared to C9-ve ALS, ALS-F and HC (p<0.05, p<0.01 and p<0.001 respectively). ROC analysis of Nf-L ICs expression distinguishes C9+ve ALS from ALS and ALS-F (**A4**, AUC=0.69; p<0.01 and AUC=0.73; p<0.01 respectively) and HC (**A4**, AUC=0.78; p<0.0001). (**A7, A8**): Nf-H Abs levels are higher in ALS-F and in the general ALS cohort compared to HC (p<0.0001 and p<0.001 respectively) and provide a modest discrimination of C9+ve ALS from HC on ROC analysis (AUC=0.65; p<0.05). (**A11, A12**): Nf-M Abs are higher in ALS-F compared to HC (**A11**, p≤0.05). (**A9, A10**): Poly (GP) DPR Abs are more expressed in C9+ve ALS patients compared to the level observed in ALS-F, ALS-S and HC (p≤0.05) and discriminate between C9+ve and the C9-ve ALS patients group (**A10**; AUC=0.72, p<0.01).

Plasma Nf-L ICs had a strong differential expression across the ALS sub-groups, with the highest level of expression in C9+ve ALS patients compared to HC (*p*=0.0002), ALS-F (*p*=0.0056) and ALS as a whole (*p*=0.040; Fig.2 A3). Of note, ALS-S had higher level of plasma Nf-L ICs expression compared to ALS-F and to HC (*p*=0.033; Fig.2 A3). Analysis of sensitivity and specificity for Nf-L Abs showed a modest effect in the discrimination of C9+ve ALS from HC (AUC=0.62; *p*=0.077) while Nf-L ICs in plasma strongly discriminated C9+ve from HC (AUC=0.78, *p*<0.0001; 84.6% sensitivity and 77.3% specificity) and from ALS-F (AUC=0.73; *p*=0.0012; 76.9% sensitivity and 73.3% specificity; Fig.2 A4). Similarly, Nf-L ICs plasma expression distinguished ALS-S from HC (AUC=0.68; *p*=0.0028) and from ALS-F (AUC=0.63, *p*=0.038; Fig.2 A4).

Nf-H and Nf-M Abs were similarly upregulated in ALS plasma samples, mainly in ALS-F patients (Fig.2 A7, *p*<0.0001 and Fig.2 A11, *p*=0.047) compared to HC. ROC analysis for Nf-H Ab levels showed a good separation of ALS from HC (Fig.2 A8, AUC= 0.68; 69.8% sensitivity and a 70.4% specificity, *p*<0.0001), ALS-F from HC (AUC=0.77; *p*<0.0001, 80% sensitivity and 77.5% specificity). ROC analysis also indicated that Nf-H Abs in plasma provided discrimination between C9+ve ALS patients and HC (AUC=0.66; *p*=0.016, Fig.2 A8). ROC curve for Nf-M Abs indicated a modest separation between ALS patients and HC (AUC=0.63, *p*=0.005; Fig.2 A12). Differences of Nf-H and Nf-M Ab levels between C9+ve ALS patients and other subgroups were not significant. Group analysis of Nf-L Abs in plasma only showed a trend towards over-expression in C9+ve patients compared to HC (data not shown).

### Poly (GP) DPRs Abs and ICs in phenotypic and genetic variants of ALS

Plasma levels of Abs and ICs to DPR were investigated in ALS subgroups, including C9orf72 mutation carriers (C9+ve) and non-mutation carriers (C9-ve). Abs against poly (GP) DPR were raised in blood from C9+ve ALS patients compared to C9-ve ALS cases, ALS-F, ALS-S and HC. Plasma levels of poly (GP) DPR Abs were significantly lower in ALS-F (*p*=0.013) and in ALS-S (*p*=0.039) compared to C9+ve patients (Fig.2 A9). ROC analysis showed that poly (GP) DPR Abs had a good performance in discriminating C9+ve from C9-ve ALS (AUC=0.72, *p*=0.0008) and C9+ve from HC (AUC=0.65; *p*=0.021, 95% CI, 0.52-0.77; Fig.2 A10). Levels of poly (GA) DPR Abs were not different among the ALS sub-groups, neither discriminated ALS from HC (data not shown).

### Survival analysis

We then evaluated whether levels of Nf protein isoforms and matched Abs and ICs affected patient survival in ALS phenotypic and genetic sub-groups. Univariate Kaplan-Meier analysis and multivariate Cox proportional hazard model adjusted for age and sex were used to assess survival and to test those factors mostly associated with ALS progression rate and risk of death. Sub-grouping of patients to test survival was based on Nf and Nf Abs/ICs expression levels, with the lower expression group being either those in the Quartile 1 (Q1, Fig.3 A1, B1, C1) or those with levels below Quartile 3 (Q1-Q3, Fig.3 A2, B2, C2). Survival analyses were performed using variables such as disease duration from either visit 1 (V1) or from symptom onset (Fig.3 A3, B3, C3). Clinical, genetic and demographic covariates such as bulbar onset, presence of a C9orf72 mutation, progression rate at V1, gender (Female), age at V1 were tested as independent predictors of survival. Kaplan-Meier analysis showed a significant association between a better survival and low levels (Q1-Q3) of Nf-H (*p*=0.0018; Fig.3 B3) and of Nf-L (*p*<0.0001; Fig.3 C3). Using Q1 as cut-off for lower expression levels, multivariate Cox proportional analysis showed an association with survival only for Nf-H (*p*=0.032; Fig.3 B1) and a trend towards significance for Nf-L protein levels (*p*=0.056; Fig.3 C1). Kaplan-Meier analysis showed association between low levels of Nf-H Abs (Q1-Q3) and survival (*p*=0.0231) while Cox multivariate analysis indicated the same association to be only close to statistical significance (*p*= 0.072; Fig.3 A3, A2 respectively).

**Figure 3.**
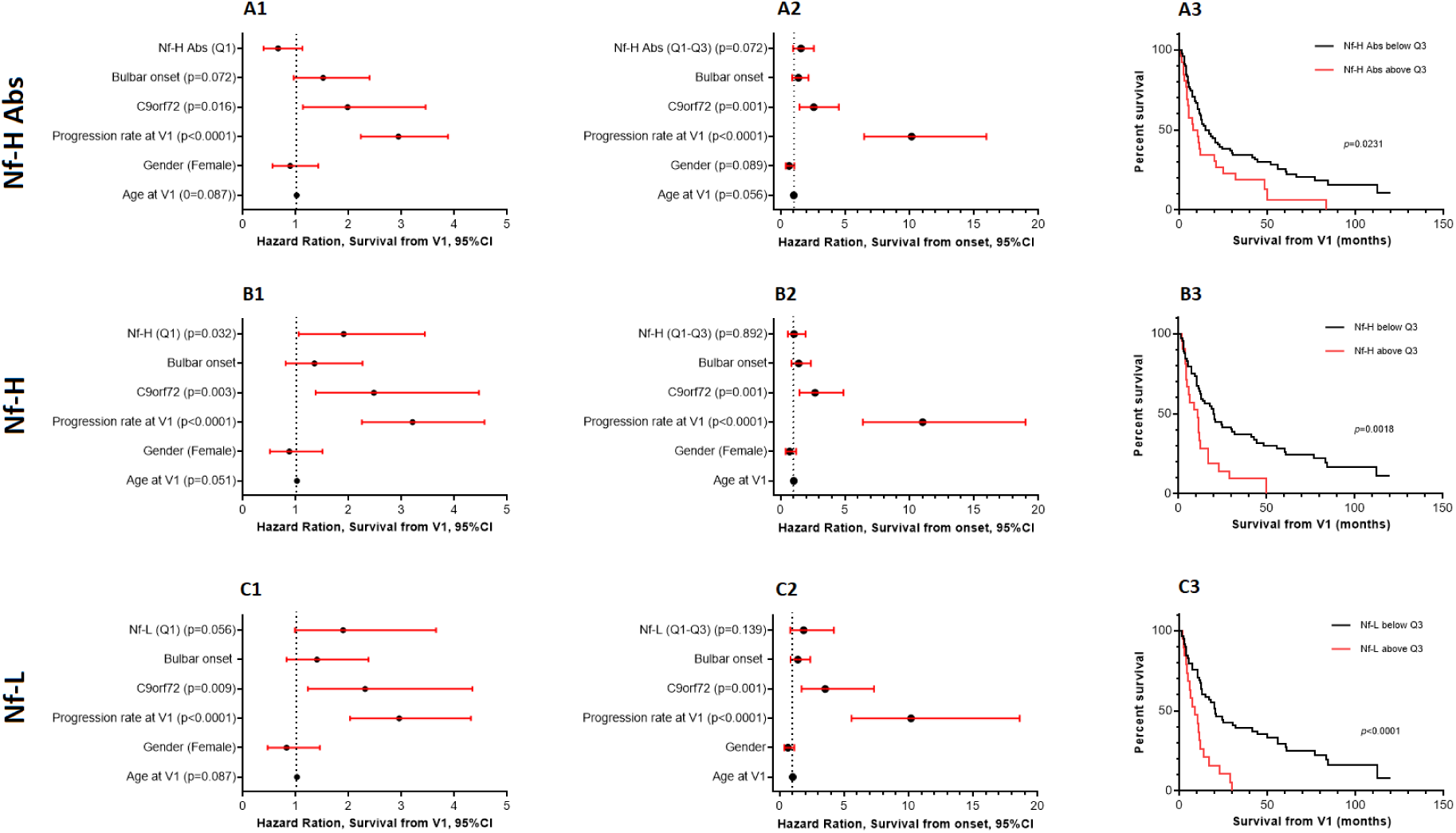
Mortality hazard ratio and Kaplan-Meier survival curves for Nf-H, Nf-L and Nf-H Abs. Mortality hazard ratio (HR) calculated by survival length. (**A1, B1, C1**): from visit 1 (V1) with sub-grouping cut-off at Quartile 1 (lower levels: Q1). (**A2, B2, C2**): from disease onset with sub-grouping cut-off at Quartile 3 (lower levels: Q1-Q3). Cox-regression analyses include covariates such as bulbar onset, C9orf72 mutation status, progression rate at V1, gender (male/female) and age at V1. Vertical dashed-line denotes unchanged mortality (HR=1). Closed circle denotes median HR and red whiskers denotes 95% CI. (**A3, B3, C3**): Kaplan-Meier survival curve from disease onset with sub-grouping cut-off at Q3. The red lines are the groups with higher level (above Q3) and black lines, groups with lower level (below Q3). *p-*value was obtained from Log-rang test, *p*≤0.05 was considered statistically significant. CI: confidence interval.

Among other variables emerging as associated with survival using Cox proportional hazard model, we have identified progression rate at V1 using both Q1 and Q1-Q3 as cut-offs for low levels of Nf-H, Nf-L and Nf-H Abs (*p*<0.0001) and the presence of a C9orf72 genetic mutation (from *p*=0.016 to *p*=0.001; Fig.3 A1, B1, C1 and A2, B2, C2).

The median patient survival for Nf-H Ab at Q1-Q3 levels was 15.75 months and above Q3 was 8.95 months (ratio: 1.76). The median patient survival for Nf-H levels at Q1-Q3 levels was 20 months and above Q3 was 10.7 months (ratio: 1.86) and for Nf-L at Q1-Q3 levels was 20 months and above-Q3 was 8.8 months (ratio: 2.2; Fig.3 A3, B3, C3).

Expression levels of Nf-M Abs, Nf-L ICs and poly (GP) DPR Abs were not good predictors of survival length (data not shown).

### Correlation between Nf isoform proteins and Nf Abs/ICs plasma expression

Spearman’s correlation analysis was also used to evaluate the strength of association between Nf proteins expression and immunoreactivity to Nf (Abs and ICs) for all ALS patients under investigation in the cross-sectional samples. There was a mild positive correlation between total Nf levels (Nf-L plus Nf-H) and total matched Nf Abs (r=0.24, *p*=0.002, Fig.4 A1) and between Nf-H isoform proteins and Nf-H Abs (r=0.18, *p*=0.02) (Fig.4 A2). A weak positive correlation was observed between total Nf and matched Nf ICs (Fig.4 B1; r=0.19, *p*=0.016) and between Nf-L and Nf-L ICs (Fig.4 B2; r=0.17, *p*=0.045).

**Figure 4.**
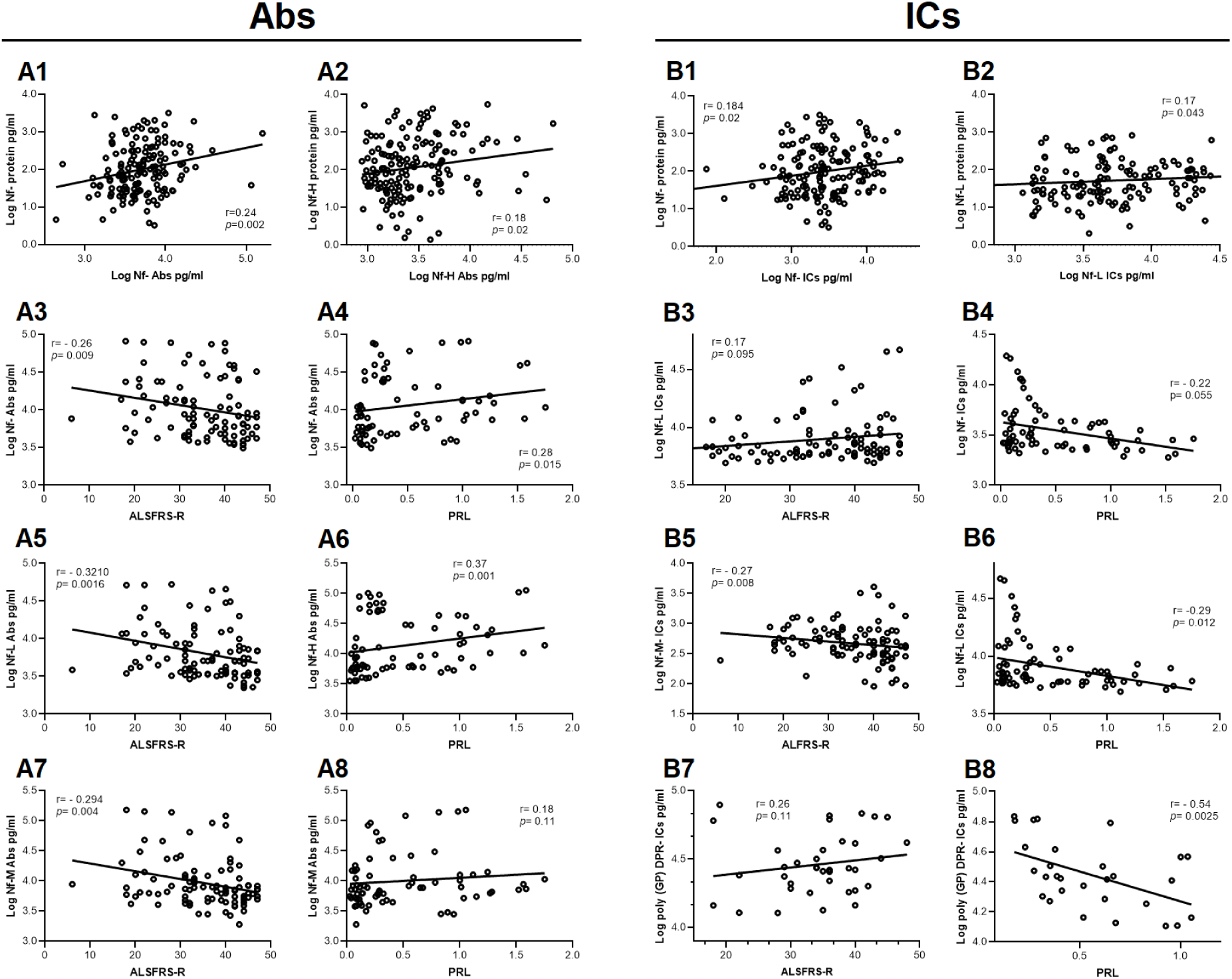
Correlation analysis between Nf Abs, ICs and functional scores representative of disease progression. Plasma samples collected from ALS patients for cross-sectional analysis (including C9-ve and C9+ve ALS patients) were tested to evaluate the relation between Nf proteins and matched Nf Abs and ICs (upper panel: **A1, A2, B1 and B2**) while plasma samples collected longitudinally were analysed to test the association between Nf Abs, Nf-L, Nf-H and Nf-M Abs / ICs as well as poly (GP) DPR-ICs with functional scores at the time of sampling (**A3 to B8**). Only significant findings (or close to significance) are shown. The upper left panels show a a positive correlation between total Nf levels and total Nf Abs (**A1**) and between Nf-H protein and Nf-H Abs (**A2**) and the upper right panels show a modest positive correlation between total Nf and matched Nf ICs (**B1**) and between Nf-L and Nf-L ICs (**B2**). Notably, different positive or negative trends were observed for the correlation analyses between Abs, ICs and ALSFRS-R and PRL for both total Nf, for Nf isoform proteins and DPR variants. A negative correlation was observed between Nf Abs and ALSFRS-R scores and a positive one with PRL (**A3, A4)**. A similar pattern was seen for Nf ICs although correlation was not significant (**B3, B4**). Notably, the negative correlation with highest significance was found to be between poly (GP) DPR ICs and PRL (*r*= −0.54, *p*=0.0025; **B8**). Spearman’s rank-order correlation was performed for the different parameters; *r=* coefficient of correlation, *p*≤0.05 was considered statistically significant. ALSFRS-R, (ALS functional ratting scale revised); PRL (progression rate at last visit).

### Correlation between Nf Abs/ICs expression and functional scores

We examined the correlation between Nf Abs and ICs plasma expression in longitudinal samples and progression rate to last visit (PRL) as well as ALSFRS-R. A negative correlation was observed between Nf Abs and ALSFRS-R scores (r=-0.26, *p*=0.009; Fig.4 A3) and a positive association with PRL (r=0.28, *p*=0.015; Fig.4 A4), while total Nf ICs showed only a trend for a positive correlation with ALSFRS-R (r= 0.17, *p*=0.095) and for a negative correlation with PRL (r= −0.22, *p*=0.055; Fig.4 B3-B4). A negative correlation was found between Nf-L Abs and ALSFRS-R (r= −0.32, *p*=0.016; Fig.4 A5) and a positive correlation between Nf-H Abs and PRL (r= 0.37, *p*=0.001; Fig.4 A6). Nf-L ICs were significantly and negatively correlated with PRL (r= −0.29, *p*=0.012; Fig.4 B6) while the most significant negative correlation was found between poly (GP) DPR ICs and PRL among C9+ve patients (*r*= −0.54, *p*=0.0025; Fig.4 B8).

### Longitudinal analysis

Changes of Nf-L, Nf-M and Nf-H Abs during disease progression are shown in Fig.5 A and B. The box plots show the ratio between the difference in Abs levels between last and baseline visits (slopes) and the time interval (in days) for both ALS-F and ALS-S. Changes over time were significantly higher in ALS-F compared to ALS-S for all Nf isoforms Abs (Nf-L Abs, *p*=0.011; Nf-M Abs *p*=0.017 and Nf-H Abs *p*=0.01; Fig.5 A). Nf-L ICs changes over time were significantly higher in C9+ve compared to C9-ve ALS patients (Fig.5 B; *p*<0.05). Time changes of all Nf Abs were also tested in ALS-F and ALS-S by Spearman’s correlation analysis of ALSFRS-R slopes (the difference of ALSFRS-R scores between later and earlier time points) and the Nf Abs (Nf-H, Nf-M and Nf-L average) slopes as reported above. A significantly negative correlation was observed in fast progressing ALS (*r*=-0.304, *p*=0.013; Fig.5 C).

**Figure 5.**
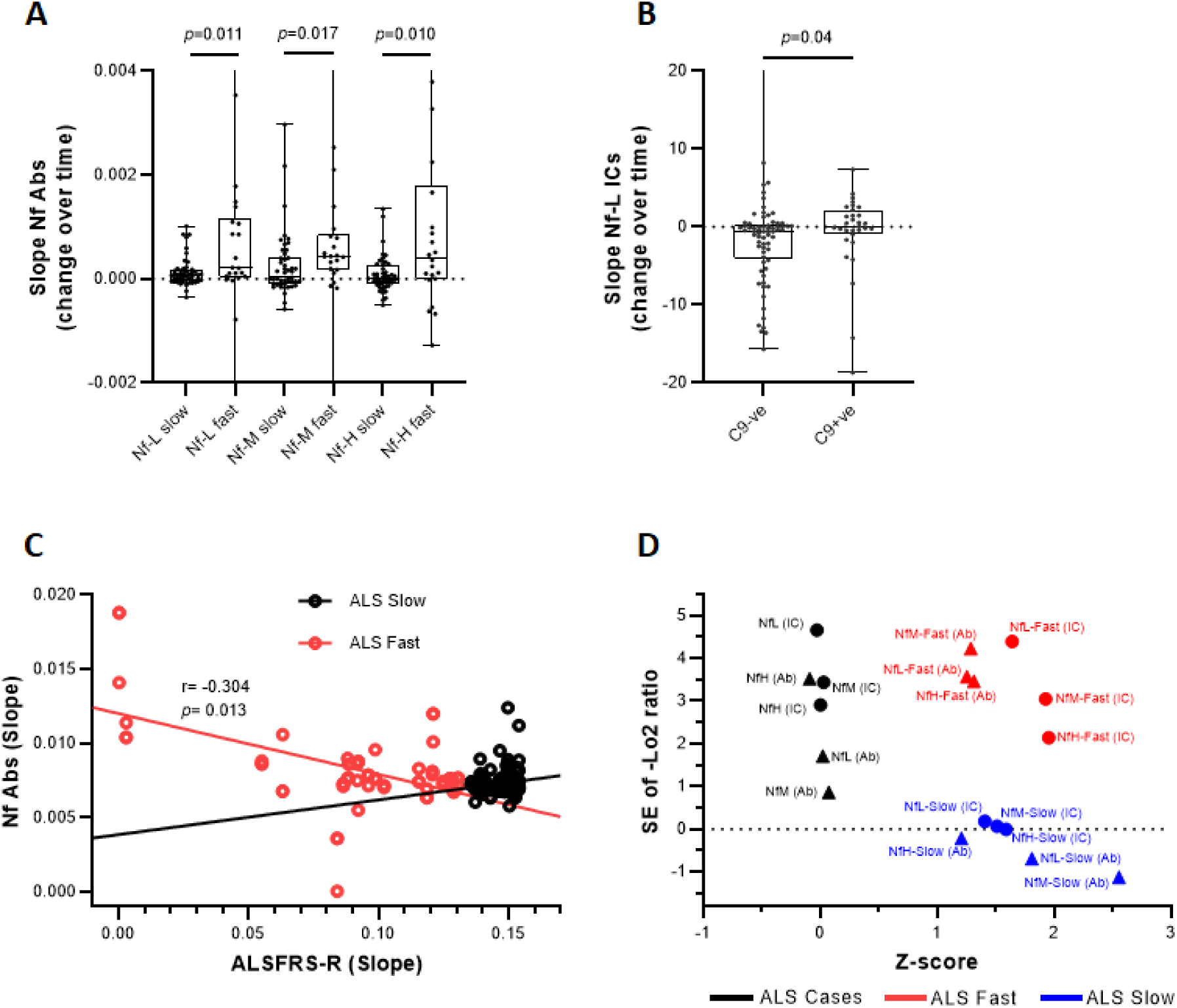
Longitudinal changes of Nf Abs and ICs in ALS. **(A):** box plots showing the change of Nf-L, Nf-M and Nf-H Abs expression levels over time. Nf Abs and ICs change is expressed as the ratio between the expression slope, calculated as the difference between last and baseline visits in fast and slow progressing ALS (ALS-F; ALS-S) over time (in days). (**B**): box plot distribution showing the changes of Nf-L ICs expression in C9+ve and C9-ve patients, expressed as slopes of expression levels between last and baseline visits over time. Box and whiskers plots show the 25^th^ and 75^th^ percentiles, median line and outliers. (**C**): changes in ALSFRS-R and Nf Abs were examined in longitudinal measurements of ALS patients with fast and slow disease progression. Spearman’s correlation was carried out between the slopes generated from changes in expression levels of Nf Abs between time points (Nf Abs slopes) and the ALSFRS-R in the same time intervals (ALSFRS-R slopes). The red line indicates a significant negative correlation in fast progressing ALS (*r*= −0.304, *p*=0.013). (**D**): The stability of measurements of different analytes in longitudinal samples from ALS fast and slow cohorts was evaluated testing 1) the change in Nf- Abs/ICs levels between last and first time points (log 2-transformed values of the standard error (SE) of tLast/t1 ratios) relative to the average change of Nf Abs (Z-score, x-axis). The scatter plot shows the cumulative Z-score for each Nf-L, Nf-M and NfH Abs and ICs in ALS-F (red symbols), ALS-S (blue symbols) and all ALS cases (black symbols).

The stability of Abs and ICs markers expression levels over time in the longitudinal samples was also evaluated interpolating 1) the log 2-transformed change in Nf Abs/ICs levels between the last (tLast) and the first (t1) measurements (tLast/t1 ratios) and 2) the variability in measurements of all Nf isoforms Ab/ICs (-log 2-transformed values of the standard error of tLast/t1 ratios) in the y-axis and the average change of Nf Abs/ICs (Zeta score) in the X-axis. The Zeta score analysis of Nf-L, Nf-M and NfH Abs and ICs showed a higher level of expression over time for ICs, particularly for Nf-L ICs, in ALS-F compared to ALS-S and to a lesser extent for Nf isoform proteins Abs (Fig.5 D).

## Discussion

The immune reactivity to proteins linked to the pathogenesis of ALS may be relevant to the immuno-pathogenesis and also as disease biomarker of this neurological condition. We have already reported a late stage increase of Nf-L Abs in ALS, while others have shown higher levels of oxSOD1 Abs in slowly progressive ALS ^11,16^. A Nf-L Abs toxic effect has recently been shown in-vitro using neuronal cell cultures while worsening of the pathological phenotype of experimental encephalomyelitis can be observed after treatment of animal models with the same Abs^10,21^. The formation of ICs/Abs to brain proteins may shift equilibrium dynamics across the blood brain barrier (BBB), reducing the concentration of proteins which may be prone to aggregation within the brain, as recently shown for the A-Beta pathological burden in brain from animal models of Alzheimer’s disease^22^ (AD). In AD patients, reduced concentration in CSF of Abeta42 peptides may also reflect the accumulation of these monomers into soluble oligomers or immuno-complexes, a process leading to a possible masking of epitopes targeted by the antibodies used in most ligand-based immunoassay^23^. Levels of natural occurring autoantibodies (NAbs) with high affinity to α-synuclein have been shown to be reduced in MSA and PD patients compared to HC, suggesting a failure of this sink mechanism under pathological conditions ^24^. Similarly, we have recently shown that Nf isoforms and particularly Nf-H are sequestered into circulating protein aggregates in plasma, a process which may also skew Nf detection by ELISA^9,20^.

We cannot infer from our data any explanation for the increase of Nf-L ICs and of other Nf Abs in the peripheral circulation of fast progressing ALS. The relationship between antigen-antibody and the changes of immune response taking place when the antigen bioavailability increases are crucial to understand Nf homeostasis in ALS and in other neurodegenerative conditions. The systemic up-regulation of Nf-L ICs we have identified in blood from ALS patients could be explained by the antigen-clearing effect that Abs against raising Nf levels exert physiologically, resulting in the reported relative stability of Nf-L/Nf-H blood expression levels throughout the symptomatic phase of ALS following the steep increase in the prodromal phase of the disease shown in asymptomatic gene mutation carriers^7,25^.

It is possible, although still to be confirmed, that Abs and ICs formed in circulation upon systemic increase of Nf or DPR proteins may accelerate BBB and spinal cord barrier (BSCB) endothelial cell damage. ICs/Abs may then gain access to the CNS and spinal cord exerting a detrimental effect on brain cells which and accelerating neurodegeneration. An altered BBB permeability has been described in ALS and it is also known to occur with ageing, which is also a major risk factor for ALS^26,27^. We have shown that the plasma proteome in late stage ALS-F contains a relatively higher level of inflammatory mediators linked to cell senescence, changes which could also affect BBB function^28^.

Whist the humoral response to Nf and DPR subtypes is increased in ALS compared to healthy control ^8,29^ (Fig.1), there is only a modest positive correlation between Nf proteins and their Abs/ICs levels in blood (Fig.4). This may reflect a variable state of immunoreactivity to changing levels of brain proteins among ALS individuals sampled at different stages of the disease. For the purpose of evaluating the potential biomarker utility of Nf Abs and ICs, we have tested clinically well-characterised but heterogeneous cohorts of ALS individuals and used Nf protein expression in the same biofluids as reference^8,20,29^. This shows unequivocally that Nf-L and Nf-H proteins, to a different degree, outperform their Abs/ICs in the separation of ALS sub-groups from HC based on rate of disease progression in line with previously reported data ^8,20,29^. However, while blood expression of Nf-L ICs, Nf-H Abs and poly (GP) DPR Abs provides a good discrimination between ALS and HC, the expression of Nf-L ICs and poly (GP) DPR Abs appears to distinguish C9+ve from the rest of ALS individuals and to increase over time in fast progressing ALS. Recent studies on the expression of poly (GP) peptides in CSF from C9+ve ALS patients and cell lines have shown that the detection of these peptides is a robust pharmacodynamic and target engagement biomarker for this sub-set of ALS patients^12^. However, CSF collection poses problems particularly when serial sampling is required to monitor treatment response in advanced ALS, when patients may be too frail and uncooperative. Unlike Nf-L and Nf-H isoform protein in blood and CSF^7,25^, poly (GP) peptide levels maintain a relatively stable expression from the pre-symptomatic to a clinically manifest stage of ALS^12^. Based on what seen in the symptomatic phase of the ALS, it will be meaningful in a future extension of this study to evaluate whether Nf-L ICs/Abs and DPR Abs may in turn increase through the pre-symptomatic and clinically manifest ALS.

While the poly (GP) DPR Abs plasma over-expression in C9+ve ALS patients can be linked to overproduction of abnormal C9orf72 translation products, the up-regulation of Nf-L ICs in C9+ve ALS cases is somehow more difficult to explain. The reported potentially enhanced state of autoimmunity that C9+ve ALS individuals may develop throughout the disease process provides some clues to explain our finding. Mutations that diminish or eliminate C9orf72 function in mice cause a severe state of autoimmunity with loss of tolerance for many nervous system autoantigens^30^. The significant down-regulation of T-regulatory cells, a prominent feature in ALS individuals with fast disease progression, is also likely to contribute to a reduced immunotolerance^4,31^. It is tempting to speculate that the increased levels of Nf Abs and of NfL ICs in ALS-F and in C9+ve cases may be the result of altered immunotolerance and enhanced autoimmunity to brain antigens which become established throughout disease progression.

This study confirms the relevance of Nf-L and Nf-H proteins as blood biomarkers for the clinical stratification of ALS. We also show the potential of ALS immunomonitoring, based on the dissection of a humoral response to Nf which is abundant and measurable cost-effectively in accessible biological fluids. Samples for Abs/ICs detection could in fact be collected remotely using newly developed methods of remote microsampling, including dry blood spots^32^ and developed alongside sub-picomolar sensitivity methods for the detection of Nf proteins. Future experiments will have to be conducted using larger longitudinal cohorts, including pre-symptomatic ALS individuals, but also focus on immunoglobulins isotypes as well as on their target affinity in relation to disease progression and severity. Additionally, it will be important to calibrate future experiments on the detection of more specific immune responses, using capture methods that include proteins that reproduce the whole range of post-translational modifications that have been described in ALS.

## Acknowledgment

We thank all participants and particularly ALS patients, their families and carers for the extraordinary support they have provided to this investigation.

The Post-Doctoral Research Assistant involved in the experimental has been supported for one year by EU2020 funding (H2020 PHC-13-2014): “Efficacy and safety of low-dose IL-2 (ld-IL-2) as a Treg enhancer for anti-neuroinflammatory therapy in newly diagnosed Amyotrophic Lateral Sclerosis (ALS) patients” (MIROCALS)”

Other past and ongoing fundings that have supported the study include:

“Blood neurofilaments in the development of neurodegeneration”

MRC Industry Case Studentship - MR/M015882/1

“Going Dry: empowering neurofilament-based biomarkers studies for disease monitoring in amyotrophic lateral sclerosis”. 17-CReA-382. ALS Association US.

We are grateful for the support of the North-Thames Local Research Network who has supported the work of research nurses and laboratory technician who have been instrumental in patient recruitment, samples collection and processing.

## Author contributions

Conception and design of the study: F.P., A.M. Acquisition and analysis of data: F.P., A.M., CH.L. Recruitment of patients and controls and data collection process: OY., A.M. Writing and revision of the manuscript: F.P., A.M., CH.L., A.K., A.N., V.L., Y.B., OY., A.I., PF. All authors evaluated and approved the manuscript.

## Potential conflicts of interest

Nothing to report.

